# Zhx2 expression in the mouse brain

**DOI:** 10.64898/2026.11.11.765907

**Authors:** Sophia A. Miracle, Morgan L. Hofmeyer, Aleksandra G. Gorelik, Ava B. Glavine, Isabella C. Conti, Sophia V. Pavlidis, Hala B. Ajjawi, Erin E. Lynch, Camron D. Bryant

## Abstract

Prescription opioids like oxycodone (**OXY**; the active component in OxyContin™) continue to drive the ongoing opioid use disorder **(OUD)** epidemic due to their widespread prescription and distribution. Genetic factors influencing pharmacokinetic and pharmacodynamic properties of opioids can have profound impacts on drug response and, in turn, influence risk for OUD. We previously used quantitative trait locus **(QTL)** mapping and gene validation to identify zinc-finger homeobox 2 **(Zhx2)** loss-of-function as a quantitative trait gene and mechanism underlying increased brain concentration of oxymorphone **(OMOR)** following systemic administration of OXY in BALB/cJ females that was linked to increased OXY reward. To test whether we could rescue the behavioral phenotype, we overexpressed Zhx2 in the brain via retro-orbital injection of a brain-specific adeno-associated virus **(AAV)**. We observed restricted Zhx2 expression (especially females), with more widespread expression in males. Consequently, we did not observe any significant effect of brain Zhx2 overexpression on OXY reward as measured via conditioned place preference **(CPP)**. However, Zhx2 overexpression did protect against OXY-induced weight loss in females following a high-dose OXY regimen. Despite the effect on weight loss, Zhx2 overexpression had no effect on spontaneous or naloxone-precipitated withdrawal behaviors. We then examined native Zhx2 protein expression in the brains of the BALB/cByJ substrain (showing wild-type levels of Zhx2 expression) via immunohistochemical co-localization and found that Zhx2 is mainly expressed in neurons located in key reward, motivational, and emotion-related regions such as the hippocampus **(HPC)**, amygdala, striatum **(Str)**, and cerebellum. To summarize, Zhx2 overexpression via retro-orbital AAV administration was quite limited, yet had significant effects on opioid-induced weight loss. Furthermore, for the first time, we provide anatomical visualization of Zhx2 expression at the protein level in neurons vs. glia throughout multiple brain regions.

## INTRODUCTION

Opioid use disorder (**OUD**) remains a major public health crisis in the United States (1). The excessive/extensive prescribing of opioids began in the late 1990s, including oxycodone (**OXY**; the active component of OxyContin™), and was followed by successive waves of overdose deaths involving prescription opioids, heroin, and fentanyl, respectively (2). Since the start of the OUD epidemic in 1999, ∼806,000 individuals have died from opioid-related deaths (3). Although prescription opioids were historically a major contributor to the opioid epidemic, only a subset of individuals transition to problematic use. OUD is known to have a substantial genetic component, with heritability estimated at ∼50-60% (4,5). Despite this, very few loci and genes have been mapped specifically for OUD.

In addition to GWAS studies for OUD, GWAS can also address opioid pharmacodynamic **(PD)** and pharmacokinetic **(PK)** phenotypes (i.e., “pharmacogenetic genes; **PGx**” – CYPs, ABC transporters, opioid receptors themselves, ion channels linked directly to opioid signaling). Genetic loci containing PK/PD genes frequently have larger effect sizes and can explain a greater proportion of the variance than disease/disorder-associated loci and therefore, pharmacogenetics of PK/PD traits (e.g., drug/metabolite levels in specific tissues), receptor/transporter/enzyme expression, etc.) and acute opioid behavioral responses can be a more tractable approach to gene identification. Several PGx genes have been shown to influence PK and PD responses to opioids. The best-characterized PK example is *CYP2D6*, in humans, which metabolizes a variety of opioids, including codeine, OXY, and tramadol, into more potent metabolites (morphine, oxymorphone **(OMOR)**, O-desmethyltramadol, respectively) (6,7). Mutations causing loss-of-function **(LoF)** or duplication of CYP2D6 alleles are associated with poor and ultrarapid metabolizer phenotypes, respectively, thus altering sensitivity to their euphoric, analgesic, and adverse effects (8,9). Transporters at tissue barriers, such as influx solute carriers [**SLC**; e.g., *SLC22A1* (organic cation transporter 1)] and efflux ATP-binding cassette [**ABC**; e.g., *ABCB1* (P-glycoprotein)], further shape drug effects by mediating brain entry and elimination of opioids, including oxycodone (10,11). For example, variants in *ABCB1* (e.g., C3435T and G2677T/A) have been associated with opioid dose requirements, plasma levels, and adverse effects (12,13). Furthermore, single nucleotide polymorphisms of *SLC22A1* (e.g., rs12208357; Arg61Cys and rs72552763; Met420del) have been implicated in impaired morphine clearance and increased likelihood of developing respiratory depression and post-operative nausea/vomiting in children (14–16). Regarding PD, the *OPRM1* A118G variant (rs1799971; Asn40Asp) has been associated with variation in opioid analgesic results (17,18) and OUD risk (5,19,20). Furthermore, its mouse equivalent, A112G, has been found to produce sex-specific effects on opioid-related behaviors (21). The use of unbiased forward genetic mapping of PK phenotypes can identify specific genes and molecular mechanisms that potentiate this risk and could inform prevention and individualized treatment (22).

OXY is a potent analgesic traditionally prescribed to alleviate moderate-to-severe pain (23). OXY binds with high affinity to the µ-opioid receptor **(MOR)**, producing analgesic and euphoric effects upon activation (24). Following administration, OXY is metabolized into an inactive metabolite, noroxycodone **(NOR)**, and an active metabolite, OMOR, which is much more potent and efficacious at the MOR compared to OXY (8) which could influence risk for OUD. In humans, metabolism of OXY to OMOR is regulated by the cytochrome P450 **(CYP)** enzyme CYP2D6. Phase I metabolism occurs primarily in the liver (9,25). O-demethylation of OXY to OMOR comprises 10% of Phase I OXY metabolism, whereas the primary Phase I metabolic pathway is N-demethylation of OXY to NOR by CYP3A4 (26,27). It is estimated that 10% of OXY is metabolized via 6-keto-reduction to α-/β-oxycodol, 10% of OXY is excreted unmetabolized, and ∼28% of the administered dose remains unexplained (7,28).

Furthermore, OXY and its metabolites undergo Phase II metabolism, where, in humans, they are glucuronidated via UGT2B7, thus producing oxycodone glucuronide (29) and the inactive oxymorphone-3-glucuronide from OMOR (30). OXY and its metabolites are then excreted primarily via the kidney, with an elimination half-life of about 3.5 to 4 hours, following oral administration (31). Because Phase I metabolism is primarily hepatic and the behavioral effects of OXY and OMOR are primarily CNS-mediated, brain OMOR concentration depends largely on liver transport and metabolism of both OXY and OMOR, systemic circulating levels, and blood-brain barrier **(BBB)** transport. In the liver, P-glycoprotein effluxes drugs into bile, thus contributing to hepatobiliary elimination (12). At the BBB, OXY undergoes active uptake via the proton-coupled organic cation **(H^+^/OC)** antiporter (32–34), whereas efflux transporters, such as P-gp, limit brain accumulation of opioids (35). Thus, brain [OMOR] depends on hepatic metabolism and clearance, systemic exposure/circulation, and influx and efflux transport at the BBB.

We previously observed increased OXY reward as measured via OXY-conditioned place preference **(OXY-CPP)** in BALB/cJ females versus BALB/cJ males and compared to near-isogenic BALBc/ByJ mice, which was correlated with increased brain [OMOR] following systemic OXY administration (36). Using gene expression **(eQTL)** and metabolite QTL analysis, and subsequently CRISPR/Cas9 gene editing, we identified a causal genetic mechanism underlying the increase in brain, namely loss-of-function in the gene zinc fingers and homeobox 2 **(Zhx2),** which codes for a transcriptional repressor protein. Zhx2 is implicated in cancer biology, including hepatocellular carcinoma, as it is most abundantly expressed in the liver (37–40). Zhx2 has also been implicated in immune system regulation, cellular development, and notably, lipid metabolism (41,42), which could be important for BBB integrity. In the BALBc substrains, BALB/cJ mice have an intronic 6 kb mouse endogenous retroviral **(MERV)** insertion near a splice site that drastically decreases Zhx2 expression, whereas BALB/cByJ mice do not and have a higher, wild-type level of Zhx2 expression at baseline (43). We constitutively modeled this loss of function via Zhx2 knockout of exon 3 and we also engineered a second mouse where we removed the MERV and restored Zhx2 expression (44). However, it remains unknown whether Zhx2 acts in the brain to influence OXY behavior. Furthermore, its distribution and cell-type specificity throughout the brain, especially at the protein level, have not been characterized.

In the present study, we first wanted to determine whether restoring brain Zhx2 expression in Zhx2 loss-of-function J mice would be sufficient to decrease the female-specific increase in OXY reward observed in BALB/cJ females with Zhx2 loss-of-function. Thus, we overexpressed brain-specific Zhx2 in BALB/cJ mice via retro-orbital administration of a brain-targeted AAV9 vector and tested for changes in OXY-CPP and expanded our behavioral phenotyping to also include spontaneous and naloxone **(NLX)**-precipitated withdrawal. We also wished to identify brain region- and cell type-specific Zhx2 expression in BALB/cByJ mice. Finally, we used immunohistochemistry **(IHC)** in naïve BALB/cByJ mice showing wild-type expression levels of Zhx2 to identify brain regions containing Zhx2 expression and to quantify its colocalization with neuronal and glial markers. Our results provide the first characterization of areas exhibiting detectable levels of Zhx2 expression in the adult brain at the protein level and indicate primarily neuronal expression. These results provide brain regions and potential circuits of interest in future studies assessing Zhx2 function in behavior.

## METHODS

### Mice

BALB/cJ mice (Strain # 000651; 16 females, 16 males) were purchased from The Jackson Laboratories (**JAX**; Bar Harbor, ME USA) for Zhx2 overexpression experiments. BALB/cJ mice harbors a 6 kb MERV insertion in Zhx2, resulting in dramatically reduced expression of this gene that can be restored via gene-edited excision of the MERV (45).

CRISPR-edited BALB/cByJ mice were engineered at JAX via excision of the only coding exon (exon 3) of Zhx2, and exon3 KO **(E3KO)** mice were bred in-house. Removing exon 3 on the BALBc/cByJ background mimics Zhx2 LoF on the BALB/cJ (44), modeling the functional effect of the naturally occurring mutation on OXY-addiction model behaviors. To maintain the line, male heterozygotes **(HET)** were backcrossed to BALB/cByJ females purchased from JAX (Strain # 001026) to produce Zhx2 E3KO HET and wild-type **(WT)** littermate mice. HET mice were then crossed to produce WT, HET, and E3KO offspring. The resulting WT female and male offspring were used for Zhx2 IHC experiments.

All mice were housed on a normal 12-hour light/dark cycle (lights on at 0630 h) and provided with food and water *ad libitum* throughout each experiment. Mice were tested at 50-120 days of age during the light phase of the light/dark cycle (0900 h-1600 h). All experiments were approved by the Institutional Animal Care and Use Committee at Northeastern University (protocol #23-1123R).

### Drugs

Oxycodone hydrochloride **(OXY)** and naloxone hydrochloride **(NLX)** were obtained from Sigma-Aldrich (St. Louis, MO, USA). Drugs were dissolved in 0.9% sterile saline and injected in a volume of 0.01 ml/g.

### Adeno-associated virus (AAV) overexpression of Zhx2 within the brain

We previously overexpressed brain-specific Zhx2 in BALB/cJ mice via intracerebroventricular administration of AAV vectors, which resulted in minimal, restricted overexpression of Zhx2 (44). Here, we attempted to obtain greater viral transfection and more widespread Zhx2 overexpression, using both a different construct (AAV9 instead of AAV/F) and a different method of injection (retro-orbital instead of intracerebroventricular).

Separate adeno-associated virus **(AAV)** viruses were designed with guidance from PackGene (Houston, TX). For systemic delivery, we used an AAV9 serotype with a CMV promoter to drive overexpression within the central nervous system (46–48). To further restrict viral expression to the brain, microRNAs miR-122 and miR-1 were included in the construct to prevent AAV viral transfection of the liver and heart, respectively (49–52). Vectors contained fluorescent eGFP reporters to validate viral placement and to facilitate IHC analysis.

Female and male BALB/cJ mice containing the 6kb Zhx2 MERV and vastly reduced Zhx2 expression (36,44) were purchased from JAX at 7 weeks of age. Mice were allowed to habituate to the colony for 1 week prior to procedures. All mice underwent retro-orbital injections for AAV administration. Mice were randomly assigned to receive an experimental AAV virus (AAV9-CMV-mZhx2-P2A-eGFP.miRT122.miRT1) to selectively overexpress Zhx2 in the brain or a control virus (AAV9-CMV-eGFP.miRT122.miRT1). Mice were heavily anesthetized with 3% isoflurane and placed in a nose cone with 2% ISO for maintenance throughout the procedure. For analgesia, 0.5% proparacaine hydrochloride (Medline; 61314-016-01) was applied to the eye and left on for ∼20 s before injection. A 29-gauge insulin needle/syringe was placed behind the right eye, and 100 ul virus was administered at a concentration of 1 × 10^11^ gc/mouse. The needle was held in place for 2 s after injection to allow virus diffusion before removal. Mice were placed on a heating pad and monitored until they regained mobility.

### Oxycodone-induced conditioned place preference

Mice were trained for drug-free and state-dependent OXY-induced conditioned place preference (**OXY-CPP**; 1.25 mg/kg, i.p.) using a 9-day procedure (53). Thirty min behavioral sessions were recorded via USB cameras (Arducam; China) and tracked via ANY-Maze Software (Stoelting; Wood Dale IL). On Day **(D)** 1, mice were tested for initial preference where they were injected with saline (**SAL**; i.p.), placed on the left-paired side of the chamber, and provided open access to both sides. On D2 and D4, mice received OXY (1.25 mg/kg, i.p.) and were confined to the OXY-paired side (right side, pointed texture floor insert). On D3 and D5 mice were administered SAL (i.p.) and confined to the SAL-paired side of the chamber (left side, smooth floor insert). On D8, all mice received SAL, were placed on the left side, and were given open access to both sides to test for state-free CPP. On D9, mice received OXY (1.25 mg/kg, i.p.) and were placed on the left side and given access to both sides of the chamber to test for state-dependent CPP. The primary measure was time spent on the drug-paired side on D1, D8, and D9.

### Spontaneous and naloxone-precipitated withdrawal in Zhx2 overexpressing mice

BALB/cJ mice with the AAV overexpression and control BALB/cJ mice with the empty vector received OXY (40 mg/kg; i.p.) at 0800 h, daily (D1-D4) and were placed in a Plexiglas cylinder (15 x 33 cm) for 30 min and then returned to their home cage. Mice received SAL (i.p.) 4 h post-OXY each day and were immediately returned to their home cage. On D5, mice were placed in the Plexiglas cylinder and recorded starting at 0700 h to assess spontaneous withdrawal (∼24 h post-OXY). Immediately following recording, mice received OXY (40 mg/kg, i.p.) and were returned to their home cage. Four h post-OXY (1200 h), mice received naloxone (1 mg/kg; i.p.) to precipitate withdrawal and were recorded for 30 min. All behaviors were recorded using a front-facing high-resolution USB camera (Arducam; China) via OBS Studio software.

### Tissue collection and perfusions

Immediately following the last behavioral session, mice were euthanized via rapid, live decapitation and brains and livers were harvested. Left brain hemispheres and liver lobes were flash-frozen in chilled 2-methylbutane. Right hemispheres and liver lobes were stored in RNAlater for ∼72 h at 4°C then transferred to −80°C until further processing. A subset of brains was perfused using 1x PBS and 4% paraformaldehyde for immunohistochemical validation of Zhx2 overexpression. Tissue was stored in RNAlater for ∼72 h at 4°C, then transferred to −80°C until further processing.

### Immunohistochemistry

For validation of Zhx2 overexpression in BALB/cJ mice, perfused brains were embedded in OCT (Thermo Fisher; 23-730-571) and sliced at 30 µm on a cryostat (Leica; CM1860). Whole brains were sliced anteriorly to posteriorly, beginning with the prefrontal cortex and ending with the cerebellum. Sections were mounted directly onto glass slides and stored at −80°C until further processing. Slides were washed 3 times in 1x PBS for 5 min each to rehydrate the tissue. A hydrophobic barrier was drawn around all tissue (Thermo Fisher; R3777). Slides were incubated in blocking solution (5% normal donkey serum, 0.2% Triton X-100, and 1× PBS) for 1 h at room temperature and then primary antibody incubation (anti-Zhx2; 1:750) at 4°C overnight. Tissue was washed 3 times in 1x PBS for 5 min each and incubated in secondary antibody (donkey anti-rabbit – AF568; 1:750) for 1 h at room temperature. Slides were then washed three times in 1x PBS for 5 min each and incubated in DAPI (1:100; Thermo Fisher; EN62248) for 5 min. Slides were cover-slipped with VectaShield mounting medium (Thermo Fisher; NC1551231) and dried at room temperature. Slides were kept in the dark at 4°C until imaging.

In BALB/cByJ mice with wild-type Zhx2 genotype (no MERV insertion) and wild-type expression levels, tissue was prepared the same as above for examining colocalization of Zhx2 with markers for neuronal and glial cell types. Slides were rehydrated in 1x PBS for 5 min. A hydrophobic barrier was drawn around all slides (Thermo Fisher; R3777) before undergoing a 1 h incubation period in blocking solution (5% normal donkey serum, 0.2% Triton X-100, 1x PBS) at room temperature, followed by a 24 h primary antibody incubation period at 4°C (anti-Zhx2; 1:750, anti-NeuN; 1:500, anti-Olig2; 1:750, anti-GFAP; 1:750, anti-Iba1; 1:750). Tissue was washed 3 times in 1x PBS for 5 min each and incubated in secondary antibody (donkey anti-rabbit – AF488, 1:750; donkey anti-guinea pig – AF488, 1:750; donkey anti-chicken – AF488, 1:750; donkey anti-goat – AF488, 1:750) for 1 h at room temperature. Slides were then washed 3 times in 1x PBS for 5 min each and incubated in DAPI (1:100; Thermo Fisher; EN62248) for 5 min. Slides were cover-slipped with VectaShield mounting medium (Thermo Fisher; NC1551231) and dried at room temperature. Slides were kept in the dark at 4°C until imaging.

Imaging was performed on a Zeiss LSM 800 confocal microscope with consistent brightness and exposure across all slides, and both sexes were processed together. All images were post-processed in ImageJ with consistent adjustments.

### Video-based withdrawal scoring

Videos were hand-scored by a trained experimenter who was blinded to treatment. Spontaneous and naloxone-precipitated withdrawal were analyzed in 5 min bins and summed to calculate the total 30 min withdrawal composite score. The primary output was the overall number of times the mice exhibited the following opioid-induced withdrawal traits: rearing, teeth chattering, crouching, grooming, jumping, backing up, paw flutters, and wet dog shakes (54–56). Total fecal boli was also quantified throughout the session.

### ImageJ quantification of Zhx2 colocalization

All fluorescent image analysis was performed in FIJI/ImageJ to quantify colocalization. All images were acquired at 20x under identical microscope conditions using a Zeiss LSM 800 confocal microscope to allow for direct comparison across conditions. Brain Zhx2 expression was detected, and fluorescence intensity quantification was assessed across the following six regions of interest **(ROIs)** for colocalization analysis: nucleus accumbens **(NAc)** shell and core, basolateral amygdala **(BLA)**, dorsal hippocampus **(dHPC)**, ventral hippocampus **(vHPC)**, and cerebellum.

For regional analysis, each image was subjected to an intensity threshold to distinguish signal from background fluorescence staining, which was held constant across all images regardless of ROI. Following thresholding, images were converted into binary masks, in which pixels above threshold were retained for further downstream analysis. Each image was first split into individual channels and subjected to background subtraction. Colocalization analysis between fluorescent channels was performed using a binary-mask overlap function for spatial overlap of Zhx2 staining with the cell type-specific marker of interest, through identification of overlapping pixels that were positive in both channels. The degree of colocalization was quantified as the proportion of overlapping signals relative to the total signal area in each channel. Quantification parameters were normalized across cell types as follows: (NeuN: 100-10000 pixels^2^; Olig2: 60-6000 pixels^2^; GFAP: 300-30000 pixels^2^; Iba1: 40-1000 pixels^2^). These ranges were selected to reflect the expected size distributions and morphologies of labeled cell populations and were applied uniformly across all images within each marker group. Replicates were obtained with n = 2-4 mice per sex per cell type.

### Statistical analysis

All statistical analyses were conducted in GraphPad Prism. We had *a priori* reason to perform separate analyses for each sex, originating from our original finding regarding a female-specific QTL for brain OMOR concentration following systemic OXY administration (36) and in a subsequent study validating a female-specific increase in brain OMOR in Zhx2 KO mice, as well as sex differences in OXY-CPP (44). Thus, for each sex, OXY-CPP was analyzed via 2-way ANOVAs (Treatment, Day) to examine the effect of treatment on distance and time spent on the OXY-paired side across D1, D8, and D9. OXY-induced weight loss was also analyzed via 2-way ANOVAs (Treatment, Day) to test for the effect of Treatment (overexpression) on weight loss following repeated OXY exposure. In cases of significant Treatment x Day interactions, Sidak multiple comparisons (padjusted<0.05) were used to identify the sources of interactions. Spontaneous and naloxone-induced precipitated withdrawal was analyzed via unpaired t-tests between treatments. Zhx2 colocalization across cell types was analyzed via one-way ANOVAs to compare overall colocalization percentages across cell types and brain regions. In cases of significant Region x Cell Type interactions, Tukey’s multiple comparisons (padjusted<0.05) were used to identify the sources of those interactions.

## RESULTS

### Zhx2 brain overexpression in BALB/cJ females and BALB/cJ males following retro-orbital AAV administration

Using retro-orbital AAV administration, we achieved a more pronounced and more widespread pattern of Zhx2 overexpression **(OE)** in BALB/cJ males vs. females (**Fig. 1A-D**). IHC analysis of males identified Zhx2 immunoreactivity in multiple limbic regions, including the fornix, BLA, dHPC, vHPC, paraventricular nucleus **(PVN)**, and periaqueductal gray **(PAG)**. IHC analysis of females showed restricted, focal Zhx2 OE in the ventral hippocampus (**Fig. 1A**). Antibody details are provided in **Supplemental Table 1**.

**Figure 1.**
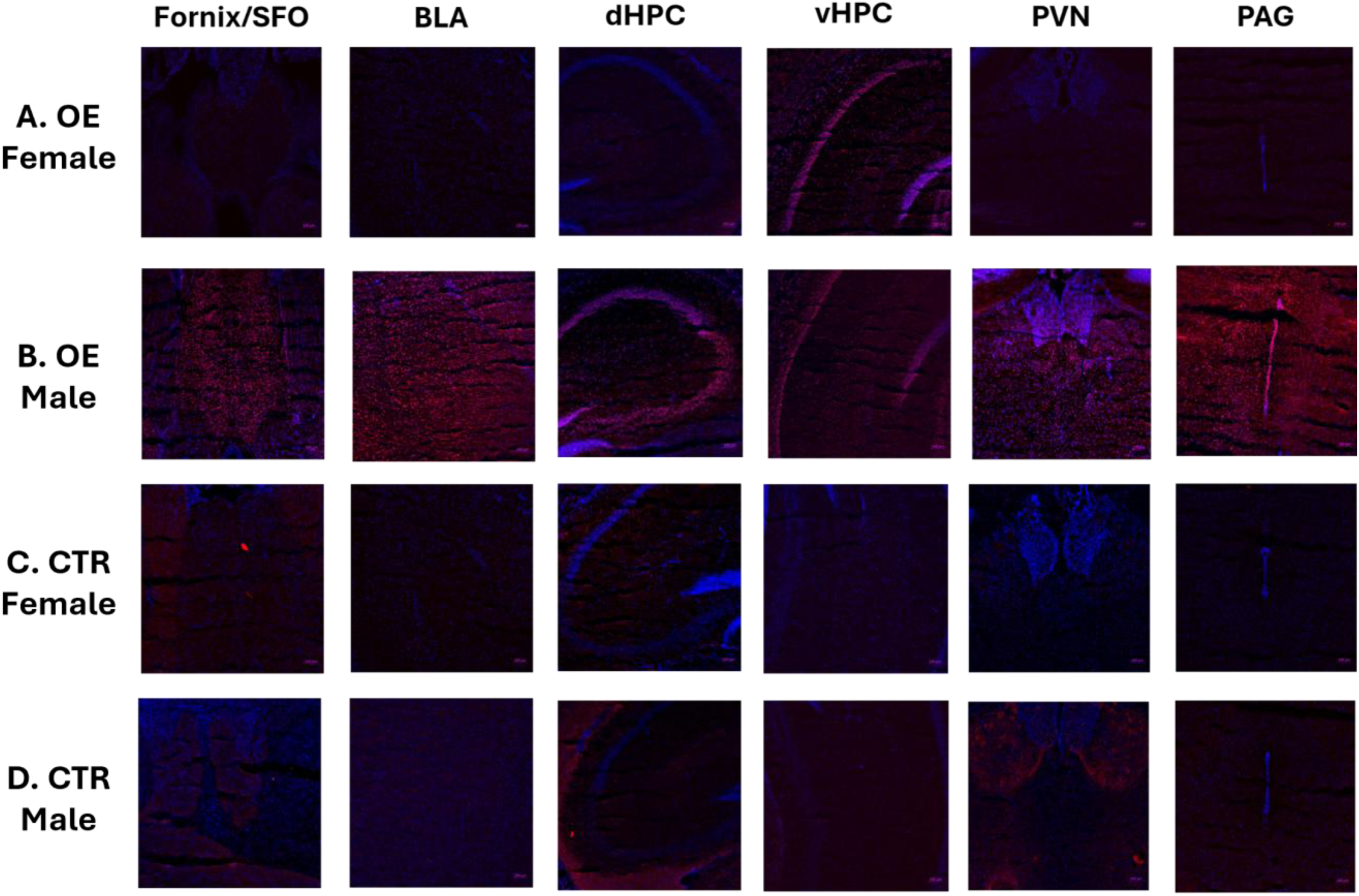
Brain Zhx2 overexpression in BALB/cJ females and males. **A-D.** Representative IHC images showing Zhx2 overexpression throughout the brain of females and males. We see robust differences in AAV transfection throughout limbic, motivation, and learning-related brain regions across sexes. Zhx2 = red; DAPI = blue; Control **(CTR)** n = 8 (4F, 4M); Zhx2 OE n = 8 (4F, 4M).

### No effect of brain Zhx2 overexpression on OXY-CPP

Previous behavioral analyses in near-isogenic BALB/c substrains revealed genetic and sex differences in OXY preference, with BALB/cJ females showing robust OXY-CPP compared to BALB/cJ males and BALB/cByJ mice that was genetically linked to reduced Zhx2 expression and increased brain OMOR concentration (36). Here, we tested the hypothesis that Zhx2 OE would result in decreased OXY-CPP (**Fig. 2A**). There was no significant effect of Zhx2 OE on either distance traveled [2-Way ANOVA: Day (F _(2, 28)_ = 24.32; p = < 0.0001); Treatment (F _(1, 14)_ = 1.397; p = 0.25); Day x OE interaction (F _(2, 28)_ = 0.89; p = 0.42)] or time spent on the OXY-paired side (2-Way ANOVA: Day [F _(2, 28)_ = 11.61; p = 0.0002); Treatment (F _(1, 14)_ = 1.291; p = 0.2750); Day x OE interaction (F _(2, 28)_ = 0.38; p = 0.69)] across days in females (**Fig. 2B, C**). Similarly, Zhx2 overexpression in males did not affect time on the OXY-paired side (2-Way ANOVA: Day (F _(2, 28)_ = 10.09; p = 0.0005); Treatment (F _(1, 14)_ = 2.88; p = 0.11); Day x OE interaction (F _(2, 28)_ = 2.18; p = 0.052)] or distance [2-Way ANOVA: Day (F _(2, 28)_ = 27.26; p = < 0.0001); Treatment (F _(1, 14)_ = 1.268; p = 0.2790); Day x OE interaction (F _(2, 28)_ = 0.43; p = 0.6562)] (**Figs. 2D, E**).

**Figure 2.**
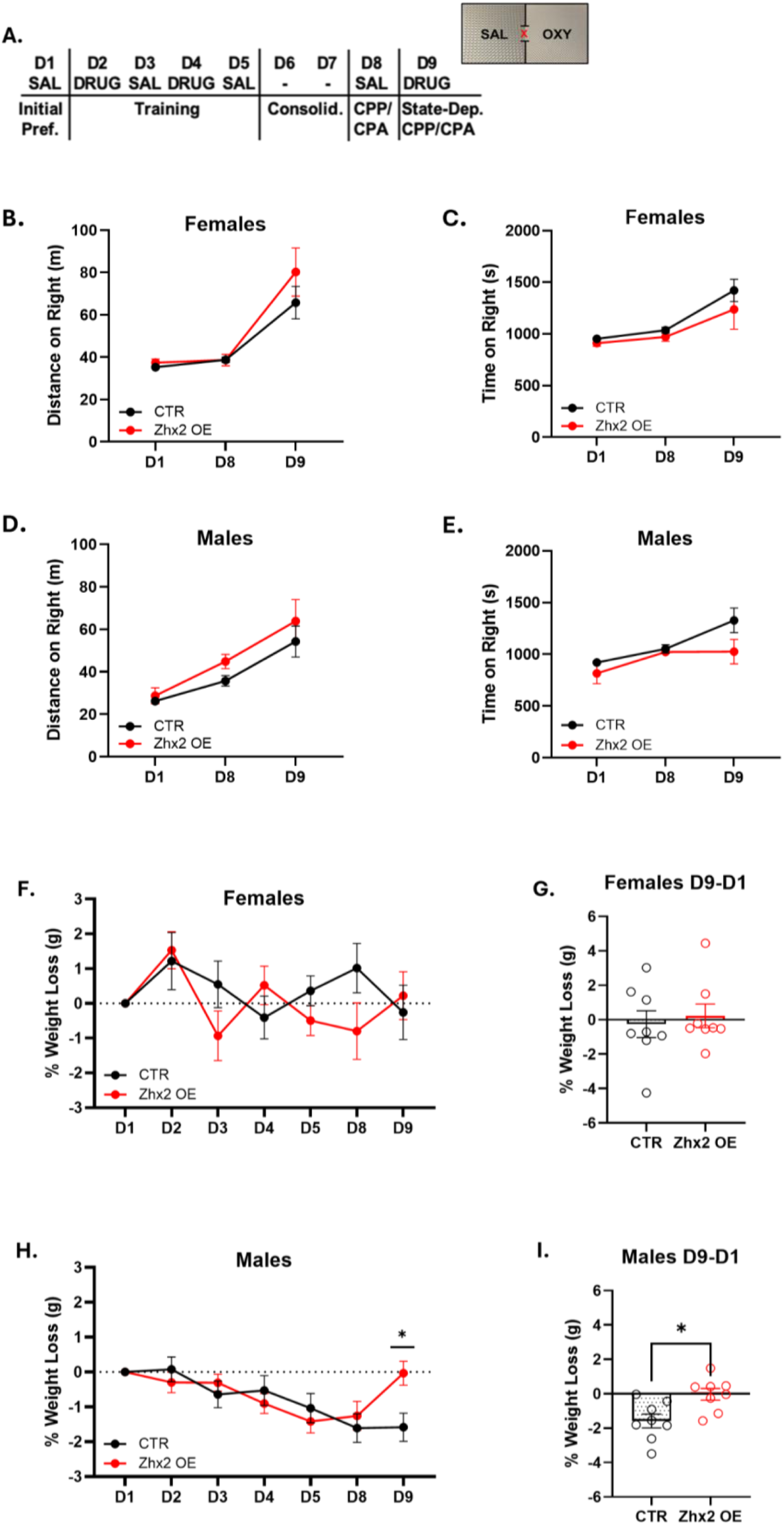
Zhx2 overexpression had no effect on OXY-CPP in BALB/cJ females or BALB/cJ males. Data are plotted as mean ± SEM. **A.** Schematic of OXY-CPP protocol. **B.** Distance traveled on OXY-paired side with overexpression **(OE)** vs. control females. **C.** Time on the OXY-paired side in BALB/cJ females. **D.** Distance traveled on the OXY-paired side in BALB/cJ males. **E.** Time spent on the OXY-paired side in males. **F.** Percent weight loss overtime in females. **G.** Percent weight loss D9-D1 in females. **H.** Percent weight loss across days in males. Zhx2 overexpression facilitated significant recovery of body weight on D9 (*p= 0.011). **I.** Percent weight loss D9-D1 in males. Control **(CTR)** n = 16 (8F, 8M); Zhx2 OE n = 16 (8F, 8M). _∗_P < .05, _∗∗_P < .01, _∗∗∗_P < .001, _∗∗∗∗_P < .0001.

In examining changes in body weight during the OXY-CPP protocol, in females, there was no effect of Zhx2 OE on body weight following low-dose (1.25 mg/kg) loss [2-Way ANOVA: Day (F _(6, 84)_ = 2.11; p = 0.14); Treatment (F _(1, 14)_ = 0.41; p = 0.5331); Day x OE interaction (F _(6, 84)_ = 2.09; p = 0.063); t-test: t_14_ = 0.4623; p = 0.65] (**Fig. 2F, G**). The lack of Day effect indicated there was no overall change in body weight in females. However, males showed significant body weight loss over days compared to females, and Zhx2 OE induced a significant recovery of this body weight loss on D9 [2-Way ANOVA: Day (F _(6, 84)_ = 6.77; p = <0.0001); Treatment (F _(1, 14)_ = 0.3063; p = 0.5887); Day x OE interaction (F _(6, 84)_ = 2.856; p = 0.0140), Šídák’s post-hoc test (D9 *p = 0.011); t-test: t_14_ = 2.937; p = 0.0108] (**Fig. 2H, I**). To summarize, Zhx2 OE had no significant effect on OXY-CPP in BALB/cJ females or males, but facilitated recovery from OXY weight loss in males that was associated with more overall weight loss in males.

### No effect of Zhx2 overexpression on spontaneous or naloxone-induced precipitated OXY withdrawal, but protection from OXY-induced weight loss in female BALB/cJ mice

We also explored the effect of Zhx2 OE on spontaneous and naloxone-precipitated OXY withdrawal (**Fig. 3A**). We hypothesized that Zhx2 OE would reduce OXY withdrawal, as Zhx2 expression is associated with reduced brain [OMOR] (36,44). There was no significant effect of Zhx2 OE on spontaneous withdrawal in either females (t-test: t_10_ = 0.9634; p = 0.36) or males (t-test: t_13_ = 1.246; p = 0.23) (**Figs. 3B, C**). There was also no significant effect of Zhx2 OE on naloxone-precipitated withdrawal in either females (t-test: t_12_ = 1.889; p = 0.08) or males (t-test: t_12_ = 0.94; p = 0.36), although there was a trending effect for reduced withdrawal in females (**Fig. 3D vs. 3E**). Interestingly, following repeated high-dose OXY (40 mg/kg) administration, Zhx2 overexpression significantly reduced the amount of spontaneous and naloxone-precipitated weight loss in females (2-Way ANOVA: Day [F _(5, 70)_ = 36.10; p = < 0.0001); Treatment (F _(1, 14)_ = 18.73; p = 0.0007); Day x Treatment interaction (F _(5, 70)_ = 3.490; p = 0.0071), Šídák’s post-hoc test (D5 *p = 0.0357) and (D5 post-naloxone **p = 0.0035)] but not in males [2-Way ANOVA: Day (F _(2.346, 32.84)_ = 45.96; p = < 0.0001); Treatment (F _(1, 14)_ = 0.1109; p = 0.74); Day x Treatment interaction (F _(5, 70)_ = 1.378; p = 0.24)] **(Fig. 3F and G)**. Analysis of individual withdrawal traits is provided in **Supplemental Fig. 3 and 4**.

**Figure 3.**
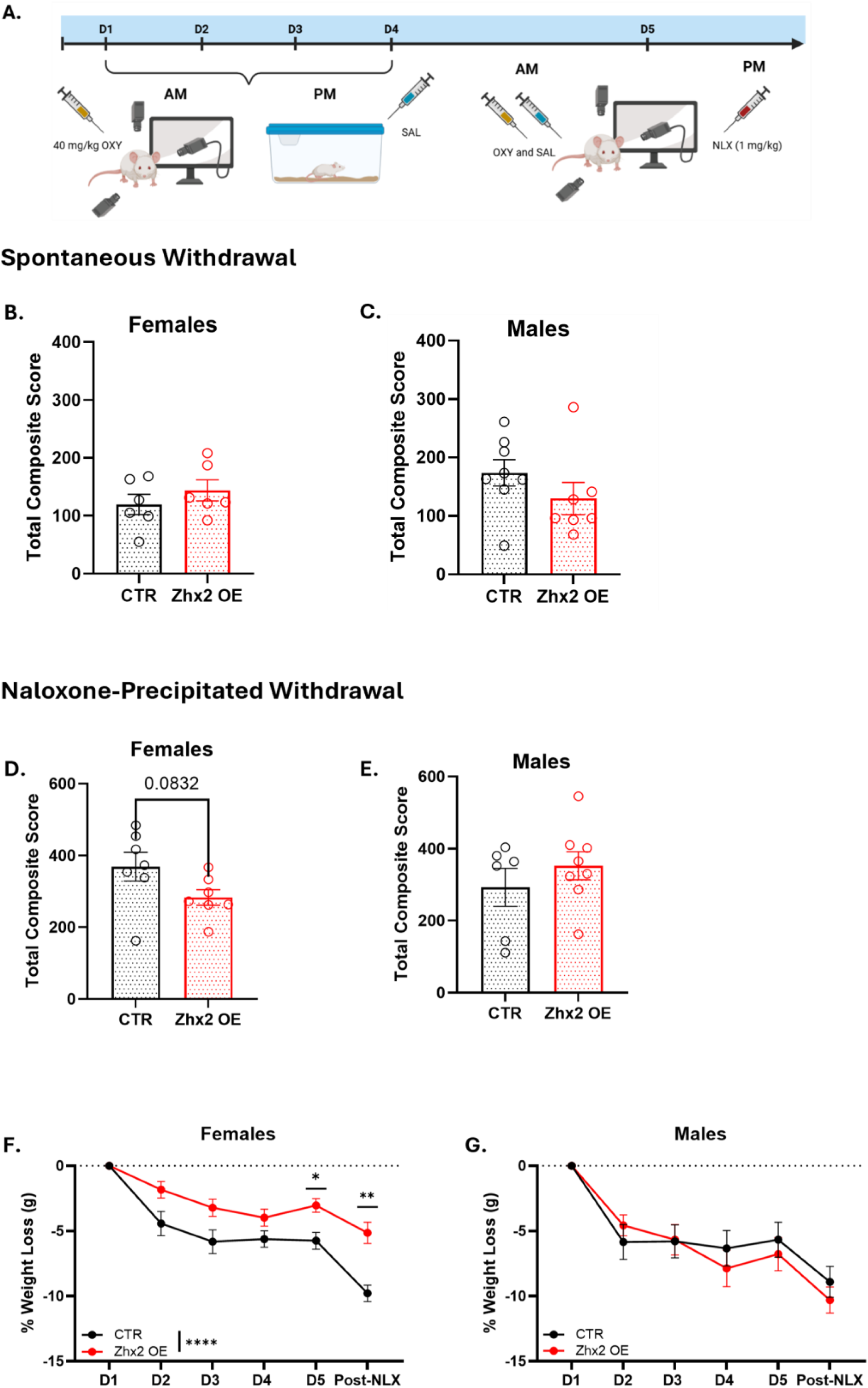
Spontaneous and naloxone-precipitated withdrawal in BALB/cJ mice with Zhx2 overexpression. Data are plotted as mean ± SEM. **A.** Schematic of naloxone **(NLX)**-induced and spontaneous withdrawal protocol. **B,C.** Composite spontaneous withdrawal score for females and males. Control **(CTR)** n = 14 (6F, 8M); Zhx2 OE n = 13 (6F, 7M). **D,E.** Composite score for NLX-induced withdrawal in females and males. CTR n = 14 (6F, 8M); Zhx2 OE n = 14 (7F, 7M). **F.** High-dose OXY-induced weight-loss in females. OE induced a significant reduction in weight loss on D5 during spontaneous withdrawal (*p = 0.0357) and also on D5 post-NLX (**p = 0.0035). **G.** High-dose OXY-induced weight-loss in males. *P < .05, _∗∗_P < .01, _∗∗∗_P < .001, _∗∗∗∗_P < .0001.

**Figure 4.**
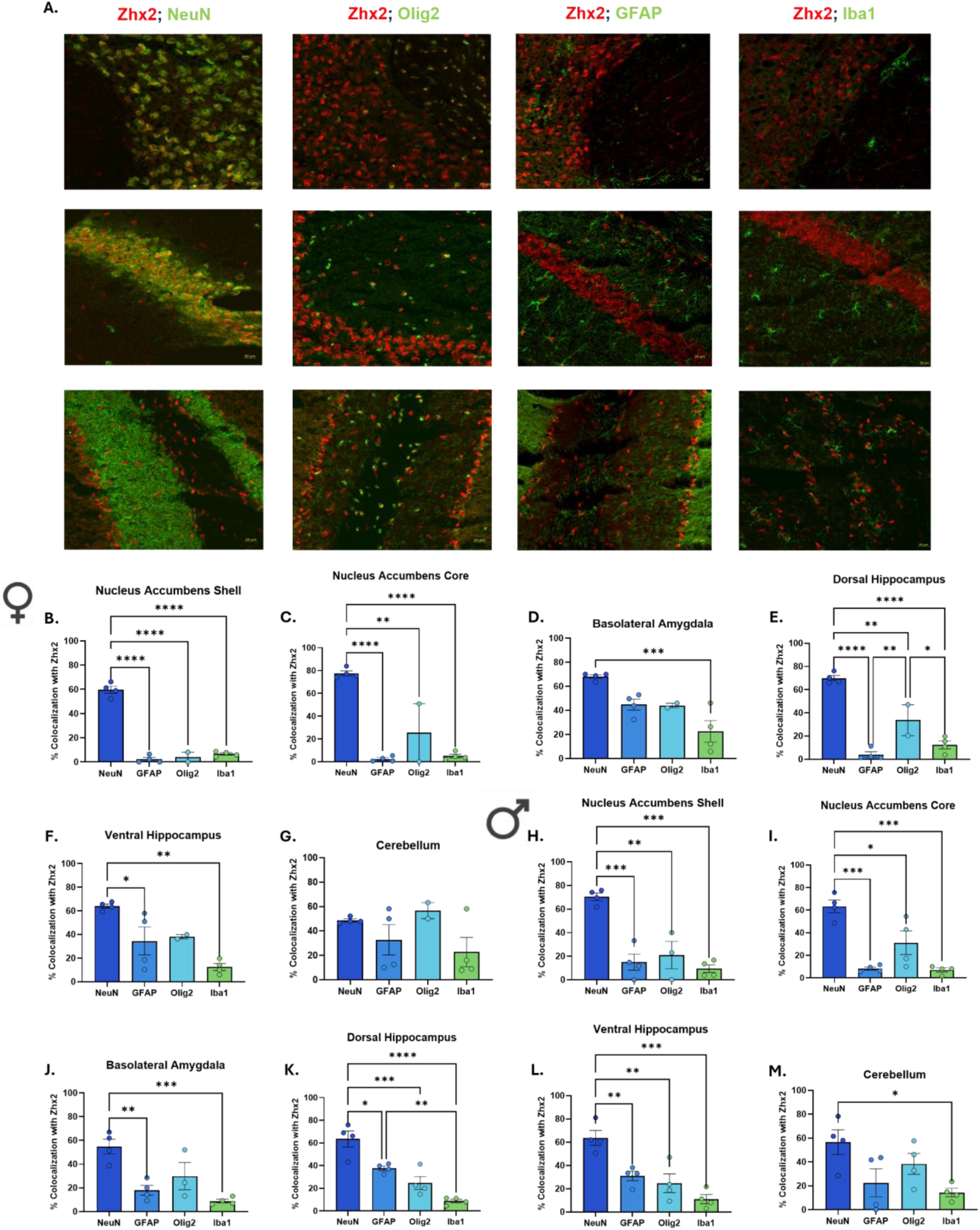
Zhx2 is highly colocalized with neurons throughout multiple regions of the BALB/cByJ mouse brain. Data are plotted as mean ± SEM. **A.** Representative 20x immunohistochemical images depicting Zhx2 (red) colocalization across cell types (green) throughout the nucleus accumbens **(NAc)** core (top), dorsal hippocampus (middle), and cerebellum (bottom). **B-G.** Females: Percent colocalization of Zhx2 across cell types in NAc shell, NAc core, BLA, dHPC, vHPC, cerebellum. **H-M.** Males: Percent colocalization of Zhx2 across cell types in NAc shell, NAc core, BLA, dHPC, vHPC, cerebellum. NeuN n = 8 (4F, 4M); GFAP n = 8 (4F, 4M); Olig2 n = 5-6 (2F, 3-4M); Iba1 n = 8 (4F, 4M). Significance was defined as follows: _∗_P < .05, _∗∗_P < .01, _∗∗∗_P < .001, _∗∗∗∗_P < .0001.

### Co-localization of Zhx2 with neurons vs. glial cell types in native wild-type BALB/cByJ mouse brain

Given the restricted effect of Zhx2 overexpression in BALB/cJ mice that otherwise have virtually zero Zhx2 expression because of the MERV insertion (45), we also wanted to examine the BALB/cByJ strain, which shows wild-type levels of Zhx2 expression, with the goal of identifying the brain regions and cell types in which Zhx2 is natively expressed. Using IHC, we assayed an extensive cell-type panel to test for co-localization of Zhx2 with neurons, oligodendrocytes, astrocytes, and microglia across multiple brain regions in which we found detectable levels of Zhx2 expression, including the NAc core, dHPC, and cerebellum (**Fig. 4A**). Zhx2 expression was significantly restricted to neurons throughout the NAc core and shell, BLA, dHPC, and vHPC, but not in the cerebellum, where colocalization with oligodendrocytes was observed (**Figs. 4B-G**; **Table 1**). Similar Zhx2 expression was observed in males, except that the males also showed significant neuronal colocalization in the cerebellum, compared to glia (**Figs. 4H-M**; **Table 2**). Antibody details can be found in **Supplemental Table 1**.

**Table 1.**
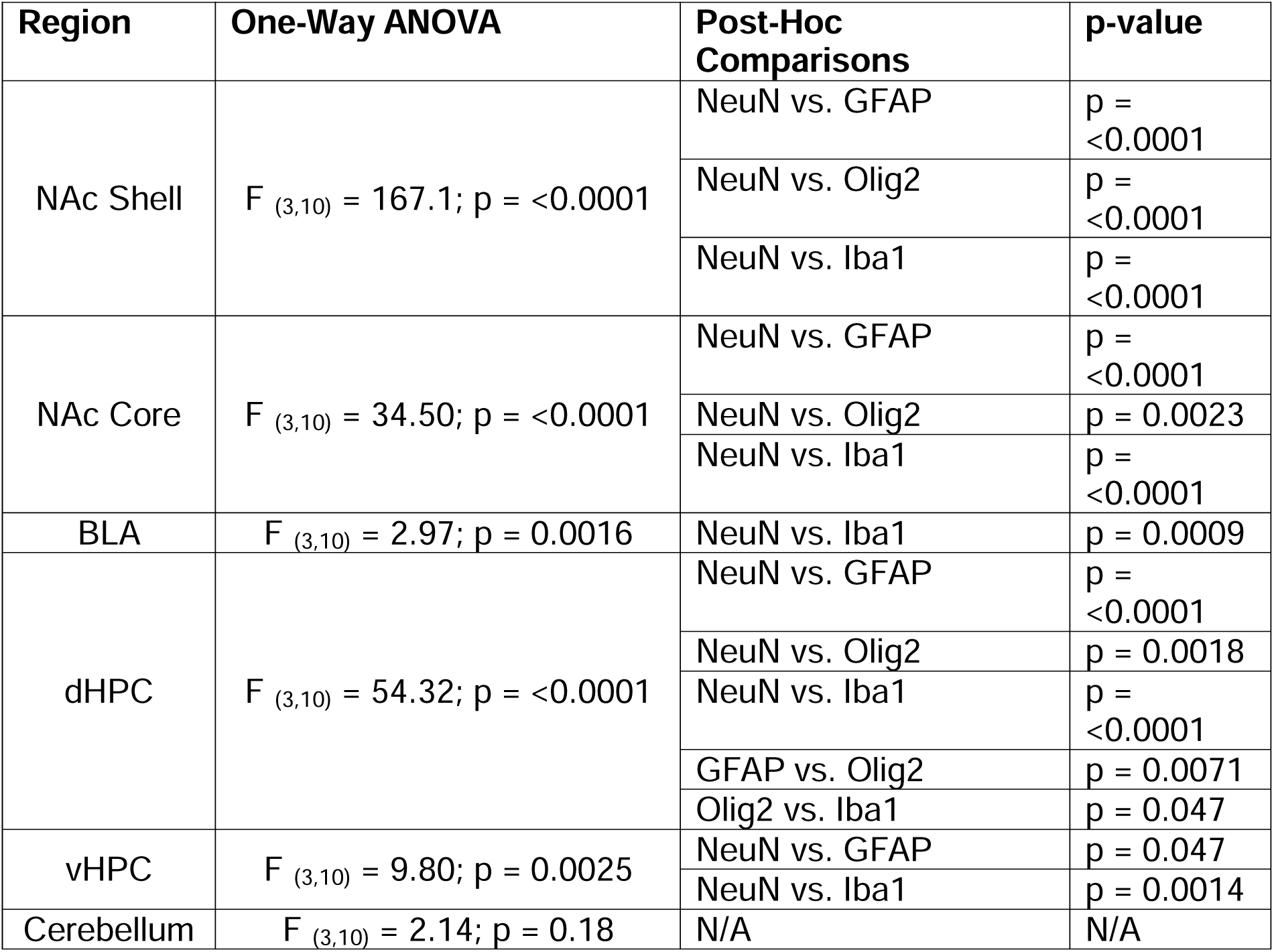
Percent colocalization of Zhx2 across cell types in female BALB/cByJ mice.

**Table 2.** Percent colocalization of Zhx2 across cell types in male BALB/cByJ mice.

| Region | One-Way ANOVA | Post-Hoc Comparisons | p-value |
|---|---|---|---|
| NAc Shell | F<sub>(3,11)</sub> = 21.09; p = <0.0001 | NeuN vs. GFAP | p = 0.0002 |
|  |  | NeuN vs. Olig2 | p = 0.0012 |
|  |  | NeuN vs. Iba1 | p = <0.0001 |
| NAc Core | F<sub>(3,12)</sub> = 18.84; p = <0.0001 | NeuN vs. GFAP | p = 0.0002 |
|  |  | NeuN vs. Olig2 | p = 0.0128 |
|  |  | NeuN vs. Iba1 | p = <0.0001 |
| BLA | F<sub>(3,11)</sub> = 11.59; p = 0.0010 | NeuN vs. GFAP | p = 0.046 |
|  |  | NeuN vs. Iba1 | p = 0.0008 |
| dHPC | F<sub>(3,12)</sub> = 23.78; p = <0.0001 | NeuN vs. GFAP | p = 0.011 |
|  |  | NeuN vs. Olig2 | p = 0.0004 |
|  |  | NeuN vs. Iba1 | p = <0.0001 |
|  |  | GFAP vs. Iba1 | p = 0.0049 |
| vHPC | F<sub>(3,12)</sub> = 14.52; p = 0.0003 | NeuN vs. GFAP | p = 0.0091 |
|  |  | NeuN vs. Olig2 | p = 0.0025 |
|  |  | NeuN vs. Iba1 | p = 0.0002 |
| Cerebellum | F<sub>(3,12)</sub> = 4.168; p = 0.0308 | NeuN vs. Iba1 | p = 0.0298 |

## DISCUSSION

We tested the hypothesis that overexpressing Zhx2 in the brain of BALBc/J mice showing loss of Zhx2 expression would reduce OXY reward and OXY withdrawal. Zhx2 overexpression was restricted to specific brain regions and had no effect on OXY locomotion or OXY-CPP in either sex. Thus, the null result on behavior could be due to the limited pattern and degree of Zhx2 overexpression. This interpretation is consistent with the fact that native Zhx2 expression in BALB/cByJ mice is more widespread in addiction circuitry (e.g., NAc, BLA, HPC) than what we observed with Zhx2 overexpression in BALB/cJ mice (A second possibility is that constitutive restoration of Zhx2 expression is required to modify the OXY behavioral profile to a BALB/cByJ-like level of wild-type Zhx2 expression – this could be addressed with the use of germline gene-edited Zhx2 overexpression. One idea related to this is that during development, Zhx2 contributes to the development of the BBB, e.g., through its role in lipid metabolism (57,58).

A third possibility is that the role of Zhx2 in OXY preference ultimately originates from a peripheral mechanism involving Zhx2, rather than the CNS. An obvious organ would be the liver, where the vast majority of OMOR is produced from OXY via CYP2D enzymes, which would then enter systemic circulation to be transported to the brain and affect OXY behavior. Our AAV construct in the present study and previous attempt at restricted CNS expression (44) included miR-122 to suppress AAV transfection in the liver. We previously were successful in robustly overexpressing Zhx2 in the liver but did not observe any significant effect on OXY behavior or brain [OMOR] (44). Thus, in this study, we were focused on potentially improving the distribution and intensity of Zhx2 expression within the brain to see if we could modify behavior. Together, the results suggest that either our CNS overexpression was insufficient to modify OXY reward or that it may require germline or early developmental rescue of Zhx2 expression within the CNS or periphery.

Zhx2 overexpression had no significant effect on spontaneous or naloxone-precipitated withdrawal, but there was a trend toward a predicted, reduced naloxone-precipitated withdrawal in overexpressing females (p = 0.08). Similar to OXY-CPP, the lack of robust effects of Zhx2 overexpression on OXY withdrawal could be due to the modest and restricted overexpression we obtained, as opioid withdrawal is mediated by multiple brain regions, including the PAG, BNST, central amygdala, and brainstem (59,60). Even though males showed more widespread AAV transfection in the PAG and BLA, there was no effect on withdrawal, suggesting that a greater level of Zhx2 overexpression is necessary that we achieved in order for a reduction in withdrawal to be observed.

Despite the lack of significant effects of Zhx2 overexpression on OXY withdrawal behaviors, Zhx2 overexpression facilitated recovery from low-dose OXY-induced weight loss during the OXY-CPP protocol in males, with no effect being observed in females, likely due to the fact that females did not show significant weight loss under these conditions.

Our immunohistochemical findings in the BALBc/ByJ strain which shows wild-type levels of Zhx2 expression revealed that Zhx2 is expressed across several brain regions implicated in reward processing, emotional regulation, and memory, including the NAc (61), BLA (62), HPC (63), and cerebellum (64). Across these regions, the percentage of Zhx2 colocalization was generally greater with neurons than with astrocytes, oligodendrocytes, or microglia, suggesting that Zhx2 function in the adult brain is limited primarily to neuronal, rather than glial, functions. These brain regions capturing the most robust Zhx2 immunoreactivity expression are relevant to opioid-related behaviors, as they contribute to reward processing and emotional and contextual memory association following drug exposure (65–67). Identification of the pattern of Zhx2 expression provides an anatomical basis for investigating site-specific Zhx2 expression in neural function underlying opioid behaviors. Of note, neuronal enrichment was observed in both sexes, and we did not identify any obvious sex differences in brain regions or cell types comprising Zhx2 expression. The Allen Brain Cell **(ABC)** atlas, which compiles snRNA-seq with multiplexed error-robust fluorescence in situ hybridization **(MERFISH)** data in C57BL/6J mice (68), indicates that Zhx2 is expressed throughout the brain, including in brain regions we identified in our study (e.g., Str and HPC). Furthermore, it corroborated widespread colocalization of Zhx2 among neuronal subpopulations throughout the brain. However, one inconsistency is that there is a robust/greater intensity of Zhx2 signal in glia, which we did not assess in this study. Perhaps the inconsistency could be explained by differences in Zhx2 RNA expression (Allen Brain Atlas) versus Zhx2 protein expression in the present study. Overall, the IHC results in the wildtype (Zhx2) BALB/cByJ strain lay the groundwork for future studies characterizing the specific neuronal subtypes expressing Zhx2 (e.g., GABAergic vs. glutamatergic).

One limitation of this study is that there was restricted AAV-mediated expression of Zhx2 throughout the brain. Following micro-injection, viral transduction is typically confined to the injection site, and the extent of diffusion depends on the serotype and its tissue-binding properties (69). AAV diffusion also depends on the host’s genetic background. In C57BL/6J mice, engineered capsids, such as AAV-PHP.eB, are able to cross the BBB more efficiently compared to BALB/c mice, which has in part been explained by strain differences in the LY6A receptor (70–72). Furthermore, strain-dependent immune responses and the ApoE-LDLR lipoprotein pathway have also been implicated (73). Because most AAV tools were optimized and validated in C57BL/6 mice, viral transduction in other backgrounds, including BALBc/J mice can be less effective. Our observations indicate that the closely related BALB/cByJ substrain shows wild-type levels of Zhx2 expression, as Zhx2 is expressed in multiple additional brain regions compared to the overexpression study in BALB/cJ mice. The more widespread Zhx2 overexpression in males vs. females could be due to the different substrain genetic background, in addition to increased neural edema as a result of Zhx2 LoF, in females (data not shown; Miracle et al., in prep). The limited Zhx2 overexpression could potentially explain the lack of effect on OXY-CPP, though any number of additional possibilities discussed above could also be responsible.

Overall, the present findings highlight sex and strain differences in AAV transfection in BALBc/J mice and suggest that focal brain Zhx2 overexpression is not sufficient to alter OXY reward or withdrawal, but it was capable of influencing OXY-induced changes in body weight in a sex- and dose-dependent manner. An important, distinct contribution of this work to the Zhx2 literature is that we describe brain regions and cell types in which Zhx2 is expressed in the wild-type BALB/cByJ strain that shows normal levels of Zhx2 expression. This documentation provides an anatomical basis for future circuit-level studies characterizing cell types and how Zhx2 expression influences OXY-addiction model behaviors.

## ACKNOWLEDGMENTS

We thank the Institute for Chemical Imaging of Living Systems (RRID:SCR_022681) at Northeastern University for consultation and imaging support.

## AUTHOR INFORMATION

### Contributions

**S.A.M.**: Experimental design, data collection, analysis, writing of manuscript; **M.L.H.**: Data collection and analysis; **A.G.G.**: Data collection and analysis; **A.B.G.**: Data collection; **I.C.C.**: Data collection; **S.V.P.**: Data collection; **H.B.A.**: Data collection; **E.E.L.**: Data Collection; **C.D.B.**: Experimental design, data collection, analysis, writing of manuscript

## ETHICS APPROVAL AND CONSENT TO PARTICIPATE

All experiments were approved by the Institutional Animal Care and Use Committee at Northeastern University.

## DATA AVAILABILITY STATEMENT

The datasets generated and analyzed during this study are available from the corresponding author upon reasonable request.

