## Supplemental Material for "Zhx2 expression in the mouse brain"

**Article Title:** Zhx2 expression in the mouse brain

**Journal:** Molecular Brain

**SUPPLEMENTAL FIGURES**

**Supplemental Figure 1. AAV plasmid maps for Zhx2 overexpression. A.** Plasmid map for the control AAV virus. **B.** Plasmid map for the experimental Zhx2-overexpressing AAV virus.

**Supplemental Figure 2. Brain regions for assessing Zhx2 colocalization with neurons and glia. A.** Nucleus accumbens core and shell (Bregma: +0.74). **B.** Dorsal hippocampus and basolateral amygdala (Bregma: -1.34). **C.** Ventral hippocampus (Bregma: -2.70). **D.** Cerebellum (Bregma: -6.24).

**Supplemental Figure 3. Spontaneous withdrawal scores for individual phenotypes.** Data are plotted as mean ± SEM. **A.** Number of paw flutters in Zhx2 overexpression **(OE)** females vs. control females. No significant difference was observed (t_10_ = 0.3080, P = 0.7644). **B.** Number of teeth chattering events in Zhx2 OE females vs. control females. No significant difference was seen (t_10_ = 1.000, P = 0.3409). **C.** Rearing events in Zhx2 OE females vs. control females. No significant difference was observed (t_10_ = 0.6627, P = 0.5225). **D.** Grooming bouts in Zhx2 OE females vs. control females. No significant difference was observed (t_10_ = 1.479, P = 0.1701). **E.** Number of backing-up events in Zhx2 OE females vs. control females. No significance was observed (t_10_ = 1.193, P = 0.2604). **F.** Number of fecal boli in Zhx2 OE females vs. control females. No significant difference was observed (t_10_ = 1.228, P = 0.2476). **G.** Number of paw flutters in Zhx2 OE males vs. control males. No significant difference was observed (t_13_ = 0.1628, P = 0.8732). **H.** Number of teeth chattering events seen in Zhx2 OE males vs. control males. No significant difference was seen (t_13_ = 0.5397, P = 0.5985). **I.** Rearing events in Zhx2 OE males vs. control males. No significant difference was observed (t_13_ = 1.288, P = 0.2202). **J.** Grooming bouts in Zhx2 OE males vs. control males. No significant difference was observed (t_13_ = 0.3084, P = 0.7627). **K.** Number of backing-up events in Zhx2 OE males vs. control males. No significance was observed (t_13_ = 0.9949, P = 0.3394). **L.** Number of fecal boli in Zhx2 OE males vs. control males. Zhx2 OE males showed significantly fewer fecal boli than control males (t_13_ = 2.210, P = 0.0456). No jumps were observed in either sex during spontaneous withdrawal. Zhx2 OE n = 13 (6F, 7M); Control n = 14 (6F, 8M). *P < .05, ∗∗P < .01, ∗∗∗P < .001, ∗∗∗∗P < .0001.

**Supplemental Figure 4. Naloxone-precipitated withdrawal scores for individual phenotypes.** Data are plotted as mean ± SEM. **A.** Number of paw flutters in control females vs. Zhx2 OE females. No significant difference was observed (t_12_ = 1.957, P = 0.0740). **B.** Number of teeth chattering events in Zhx2 OE females vs. control females. No significant difference was observed (t_12_ = 0.9234, P = 0.3740). **C.** Crouching events in Zhx2 OE females vs. control females. No significant difference was observed (t_12_ = 1.771, P = 0.1019). **D.** Rearing events seen in Zhx2 OE females vs. control females. No significant difference was observed (t_12_ = 1.637, P = 0.1275).  **E.** Grooming bouts in Zhx2 OE females vs. control females. No significant difference was observed (t_12_ = 0.3247, P = 0.7510).  **F.** Number of jumps in Zhx2 OE females vs. control females. No significant difference was observed (t_12_ = 0.4182, P = 0.6832). **G.** Number of backing-up events in Zhx2 OE females vs. control females. No significance was observed (t_12_ = 0.1846, P = 0.8566). **H.** Number of fecal boli seen in Zhx2 OE females vs. control females. No significant difference was observed (t_12_ = 1.050, P = 0.3145). No wet dog shakes were observed in females. **I.** Number of paw flutters in Zhx2 OE males vs. control males. No significant difference was observed (t_12_ = 1.960, P = 0.0736). **J.** Number of teeth chattering events in Zhx2 OE males vs. control males. No significant difference was observed (t_12_ = 1.132, P = 0.2799). **K.** Crouching events seen in Zhx2 OE males vs. control males. No significant difference was observed (t_12_ = 0.4502, P = 0.6606). **L.** Rearing events seen in Zhx2 OE males vs. control males. No significant difference was observed (t_12_ = 0.3056, P = 0.7652). **M.** Grooming bouts in Zhx2 OE males vs. control males. No significant difference was observed (t_12_ = 0.9214, P = 0.3750). **N.** Number of jumps in Zhx2 OE males vs. control males. No significant difference was observed (t_12_ = 0.9258, P = 0.3728). **O.** Number of backing-up events in Zhx2 OE males vs. control males. No significance was observed (t_12_ = 1.877, P = 0.0851). **P.** Number of fecal boli in Zhx2 OE males vs. control males. No significant difference was observed (t_12_ = 0.495, P = 0.9614). **Q.** Number of wet dog shakes events in Zhx2 OE males vs. control males. No significant difference was observed (t_12_ = 1.604, P = 0.1348). Control n = 13 (7F, 6M); Zhx2 OE n = 15 (7F, 8M). *P < .05, ∗∗P < .01, ∗∗∗P < .001, ∗∗∗∗P < .0001.

**Supplemental Table 1. Antibodies used for immunohistochemistry**

**Supplemental Figure 1.**

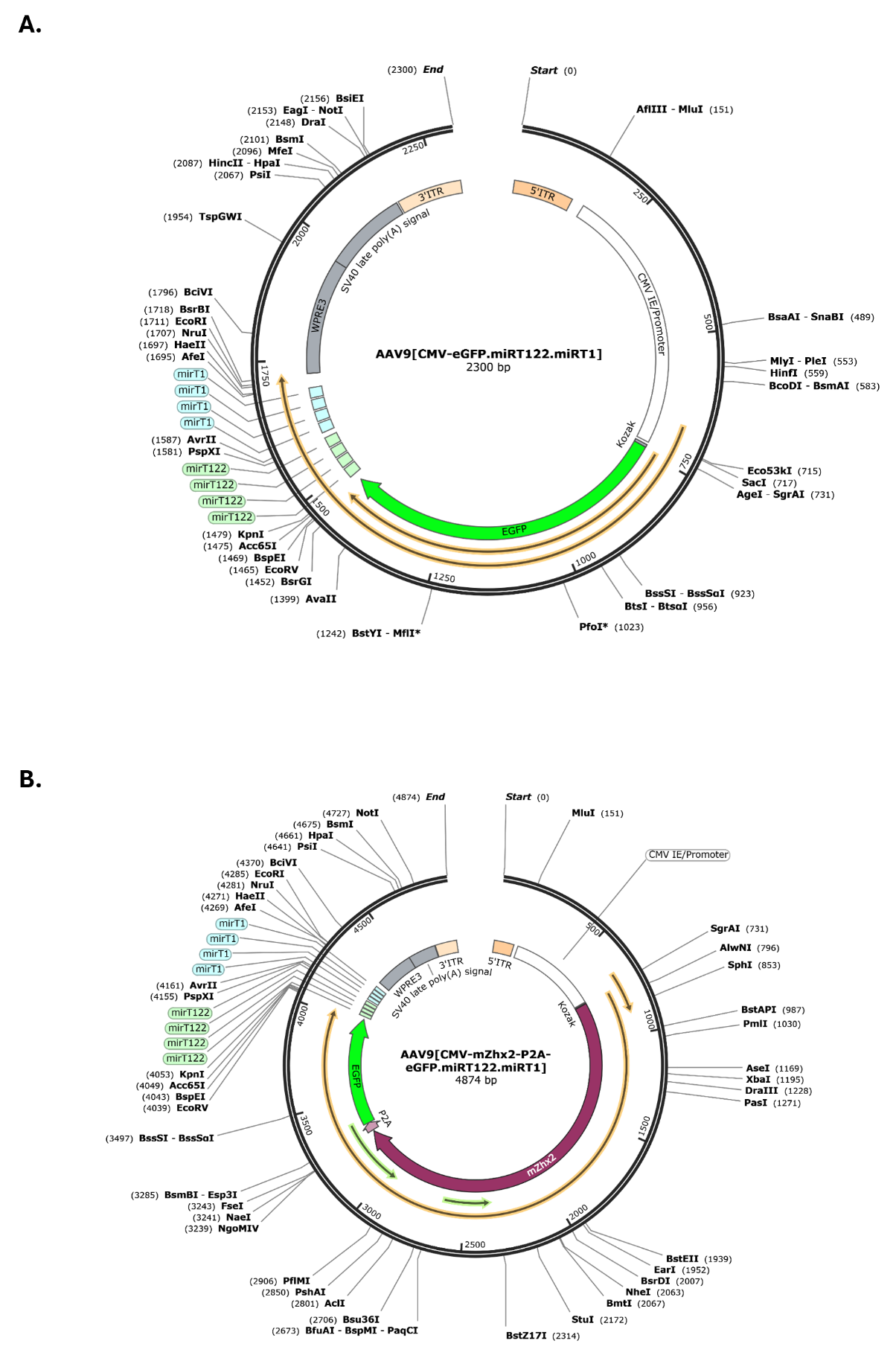

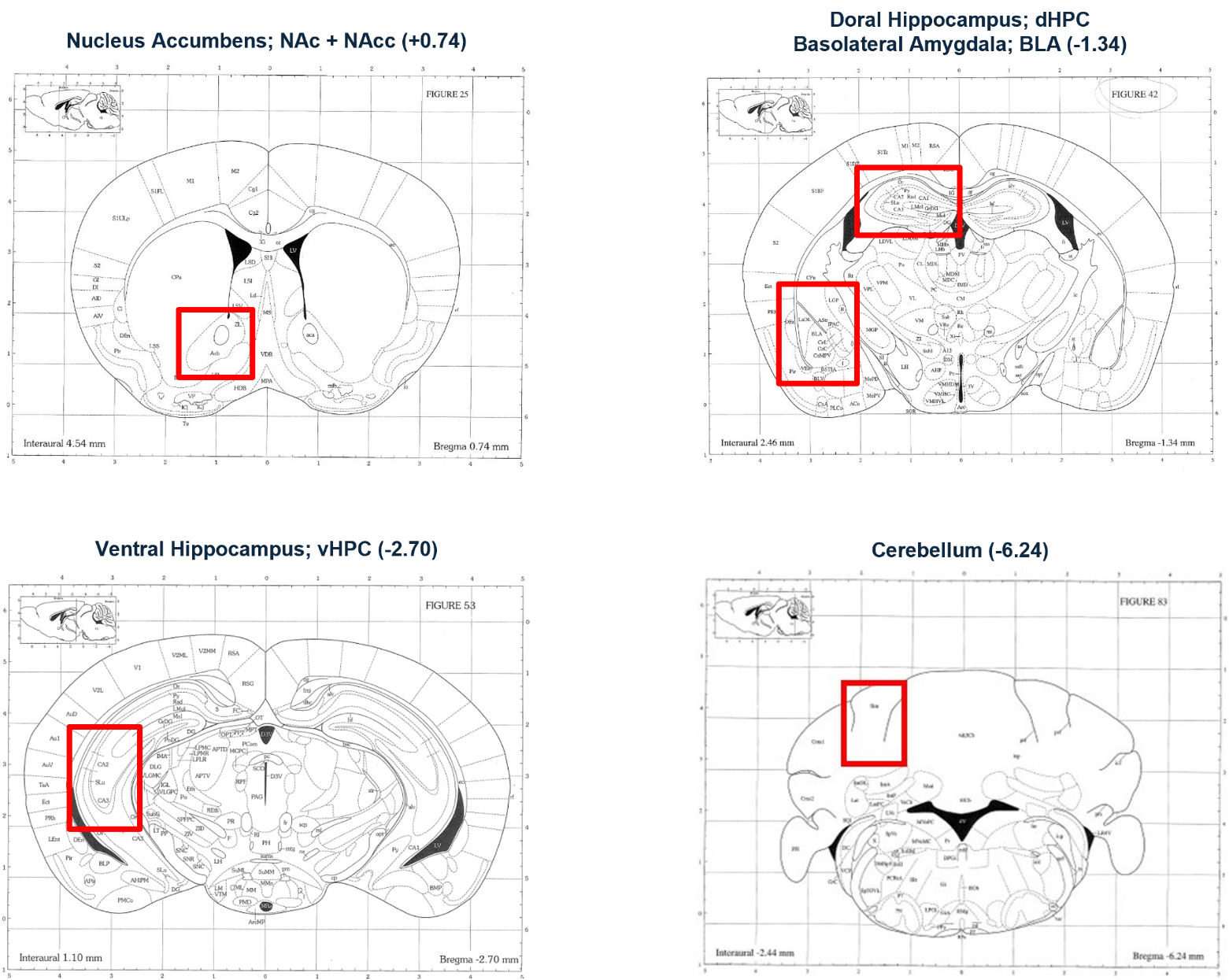
**Supplemental Figure 2.**

**Supplemental Figure 3.**

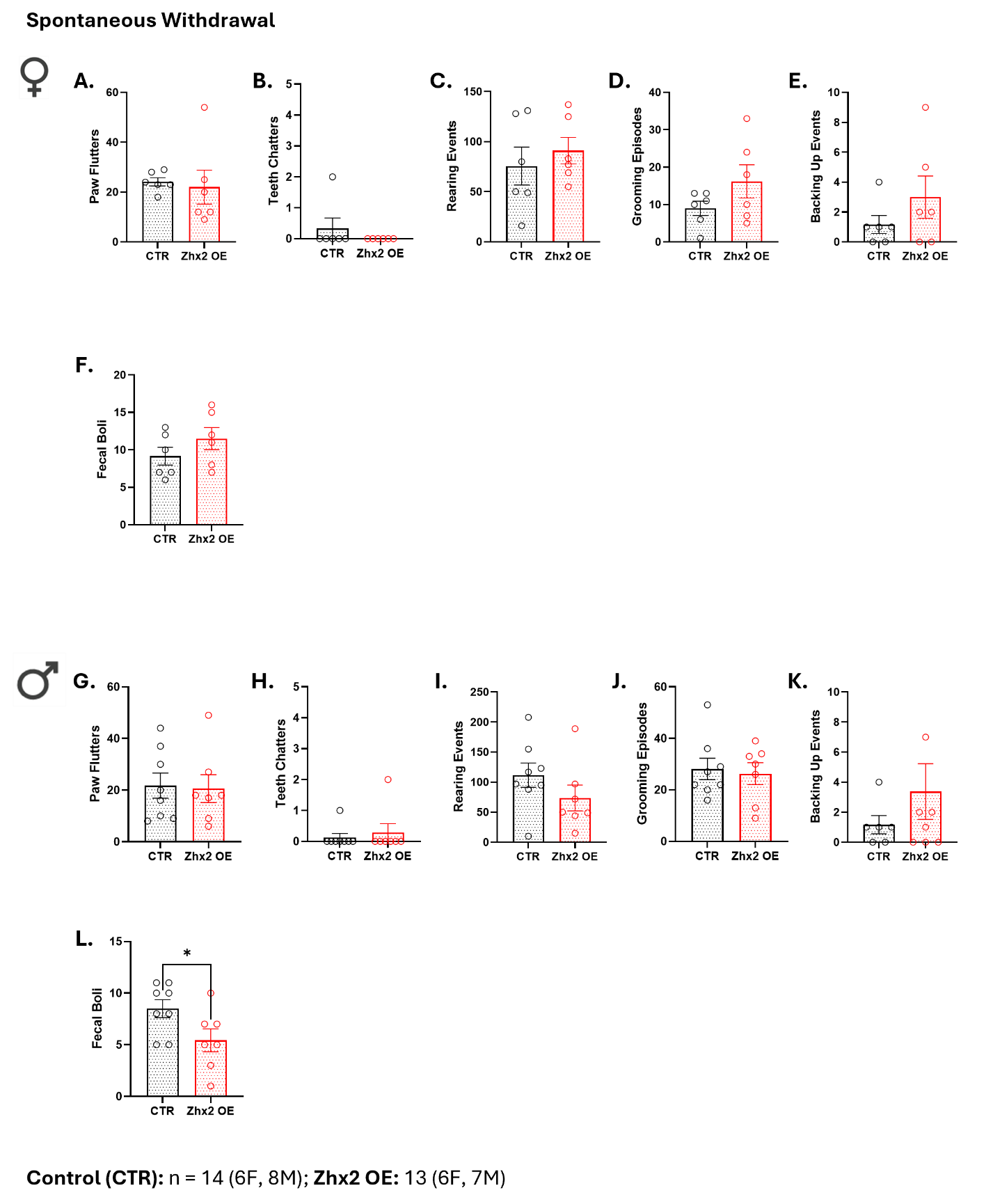

**Supplemental Figure 4.**

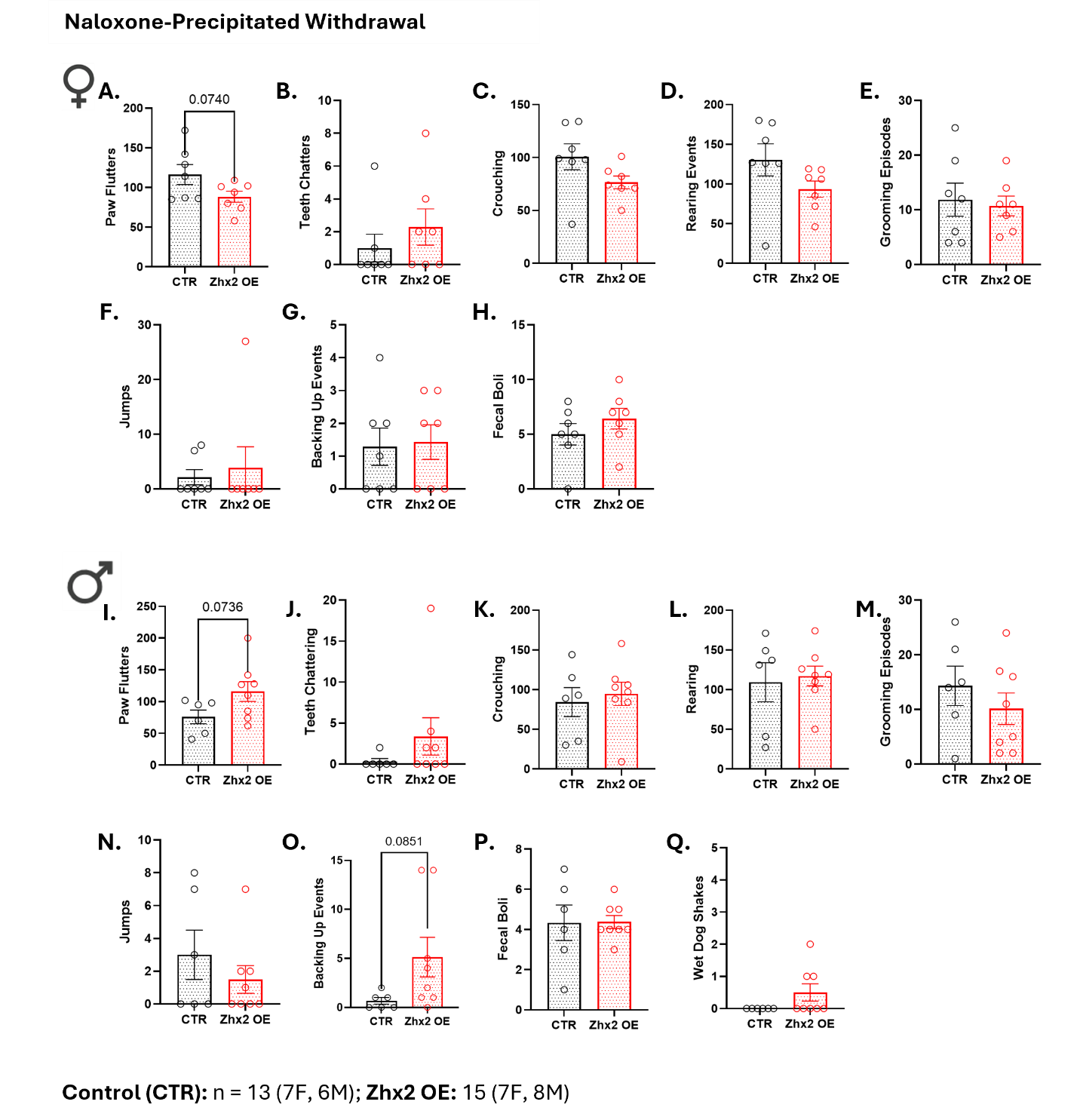

**Supplemental Table 1.**

| **Target** | **Host Species** | **Dilution** | **Alexa Fluorophore** | **Catalog #** |
| --- | --- | --- | --- | --- |
| Anti-Zhx2 | Rabbit | 1:750 | N/A | GTX112232 |
| Anti-NeuN | Guinea Pig | 1:500 | N/A | ABN90 |
| Anti-GFAP | Chicken | 1:750 | N/A | NBP105198 |
| Anti-Olig2 | Goat | 1:500 | N/A | AF2418 |
| Anti-Iba1 | Goat | 1:750 | N/A | PA518039 |
| Anti-Rabbit | Donkey | 1:750 | 568 | A10042 |
| Anti-Guinea Pig | Donkey | 1:750 | 488 | 706545148 |
| Anti-Chicken | Donkey | 1:750 | 488 | A78948 |
| Anti-Goat | Donkey | 1:750 | 488 | A11055 |
